# Cryo-EM uncovers a sequential mechanism for RNA polymerase I pausing and stalling at abasic DNA lesions

**DOI:** 10.1101/2024.09.09.611646

**Authors:** Alicia Santos-Aledo, Adrián Plaza-Pegueroles, Marta Sanz-Murillo, Federico M. Ruiz, Jun Xu, David Gil-Carton, Dong Wang, Carlos Fernández-Tornero

## Abstract

RNA polymerase I (Pol I) transcribes ribosomal DNA (rDNA) to produce the rRNA precursor, which accounts for up to 60% of the total transcriptional activity in growing cells. Pol I monitors rDNA integrity and influences cell survival, but little is known about how this enzyme processes abasic DNA lesions. Here, we report electron cryo-microscopy (cryo-EM) structures of Pol I at different stages of stalling at abasic sites, supported by in vitro transcription studies. Our results show that templating abasic sites can slow nucleotide addition by base sandwiching between the RNA 3’-end and the Pol I bridge helix. However, the presence of a templating abasic site induces opening of the Pol I cleft for either enzyme dissociation from DNA or for access of A12-Ct into the active site to stimulate RNA cleavage. Nucleotide addition opposite the lesion induces an early translocation intermediate that is different from previously-described RNA polymerase paused states, as DNA bases in the hybrid tilt to form hydrogen bonds with the newly-added RNA base. While in this state nucleotide addition is strongly disfavoured, intrinsic Pol I RNA cleavage activity acts as a failsafe mechanism to minimize lesion bypass. Our results uncover a two-step mechanism leading to persistent Pol I stalling after nucleotide addition opposite Ap sites, which is distinct from arrest by CPD lesions and from Pol II blockage at Ap sites.

## Introduction

Genome integrity is critical for cell homeostasis, an equilibrium that is challenged by DNA damage. Among the >50,000 DNA lesions per day produced in a human cell, the most abundant are abasic (Ap) sites. Around 10,000 apurinic sites are formed daily in a cell under physiological conditions, in contrast to only 500 apyrimidinic sites^1^. Ap sites arise either from spontaneous depurination or from defects in base excision repair (BER) of oxidated or alkylated bases^2^. Cells mainly repair Ap sites through additional activation of the BER pathway, while nucleotide excision repair (NER) operates as a parallel mechanism. Repair of these DNA lesions protects both from mutations in the genetic information due to loss of the base code and from genome instability due to interference with replication or transcription.

Ap sites block replication by interfering with the progression of replicative DNA polymerases (DNAPs). To overcome the obstruction, replicative DNAPs can be replaced by translesion DNAPs, which have developed different strategies to handle Ap sites. Most DNAPs follow the so-called ‘A-rule’ or ‘purine rule’ because of their preference for adenine incorporation, or guanine in a lesser degree, opposite Ap sites^3^, essentially due to the higher base-stacking ability of purines^4,5^. Alternative translesion DNA synthesis mechanisms include cytosine addition opposite a templating arginine that occupies the gap left by the missing base, as well as generation of a –1 frameshift due to Ap site extrusion from the active center^6–8^.

Besides replication, Ap sites in the DNA template strand also induce stalling of DNA-dependent RNA polymerases (RNAP). Three types of RNAPs transcribe the nuclear genome in eukaryotic cells. RNA polymerase I (Pol I) produces the ribosomal RNA (rRNA) precursor that after maturation leads to the major RNA components of the ribosome, RNA polymerase II (Pol II) mainly transcribes messenger RNA (mRNA) genes, and RNA polymerase III (Pol III) synthesizes short, untranslated RNAs such as transfer RNA (tRNA). Previous studies of Ap site management by Pol II show initial enzyme stalling at the lesion, followed by slow bypass obeying the A-rule^9^.

The nucleotide addition cycle has been best characterized in Pol II. A DNA nucleotide in the templating position (*i+1*) faces an empty active center^10,11^, where three aspartates coordinate a Mg^2+^ ion (Mg A). Arrival of a nucleoside triphosphate (NTP) opposite the templating nucleotide, in the so-called addition site (A site), promotes closure of the trigger loop to sense base paring between the incoming and templating nucleotides^12^. At this stage, the NTP phosphates, which coordinate a second Mg^2+^ ion (Mg B), are positioned so that RNA nucleotide addition to the 3’-end of the nascent RNA is catalyzed, with release of pyrophosphate and Mg B^12,13^. This provokes the backward movement of the trigger loop, which leads to enzyme translocation by one nucleotide due to conformational rearrangements of both the bridge helix and trigger loop^14^. As a result, the newly-formed base pair occupies position *i–1* in the DNA/RNA hybrid, while the next DNA nucleotide locates at *i+1* to start a new addition cycle. Apart from the A site for nucleotide addition, Pol II harbors an entry site (E site) where NTPs bind in an inverted position respect to the A site^15^. Interestingly, the A and E sites in Pol II overlap partially and, thus, cannot be occupied simultaneously. While only the cognate NTP can occupy the A site, any NTP can enter the E site, suggesting an NTP-probing role for this site. Closure of the trigger loop during nucleotide addition prevents an NTP in the A site from returning to the E site due to steric hindrance.

RNAPs are known to pause or reduce the elongation rate in response to different signals, including misincorporations in the growing RNA that alter the DNA/RNA hybrid^16^. Sensing of a distorted hybrid induces reverse translocation of the enzyme, i.e. backtracking, which inactivates nucleotide addition because the RNA 3’-end is moved away from the active center^17^. Reactivation of the enzyme from a backtracked state requires RNA cleavage, which in Pol II is stimulated by binding of the C-terminal zinc ribbon (ZnR) from transcription factor IIS (TFIIS) to the enzyme funnel^13,17^.

In growing eukaryotic cells, most transcriptional activity is devoted to rRNA synthesis and, consequently, the Pol I function is a major regulator of cell homeostasis^18^. Among the 14 subunits constituting Pol I in *Saccharomyces cerevisiae* (13 in most species) the two largest, A190 and A135, form the DNA binding cleft and harbor the active center^19,20^. Assembly of these subunits is stabilized by the AC40/AC19 heterodimer, which is shared with Pol III while peripheral subunits Rpb5, Rpb6, Rpb8, Rpb10 and Rpb12 are shared among the three nuclear RNAPs. Subunits A43 and A14 form the stalk that interacts with Rrn3^21–23^, an activating factor that binds Pol I prior to promoter recruitment^24,25^. Subunit A12 is composed by two ZnR that share homology with Pol II subunit Rpb9 and TFIIS^26^. While the N-terminal ZnR (A12-Nt) stimulates elongation^27,28^, the C-terminal ZnR (A12-Ct) is involved in RNA cleavage and proofreading^29,30^. The A49/A34.5 heterodimer, with domains that present homology with TFIIF and TFIIE in Pol II, stimulates backtracking and RNA cleavage^30,31^. This heterodimer can reversibly dissociate from Pol I to generate Pol I*^32^, thus uncovering a binding site for A12-Ct that is distant from the active center^33^.

Previous work on Pol I stalling by DNA lesions restricts to cyclobutene pyrimidine dimers (CPD), a common bulky DNA lesion induced by UV light^34^. These studies showed that Pol I and Pol II handle bulky lesions differently, with Pol I stalling firmly before CPD can access the active center while Pol II is able to slowly incorporate nucleotides opposite the lesion. To evaluate if this differential behavior is specific for bulky lesions or may also operate for other lesions, we studied the impact of Ap sites on Pol I activity.

We present biochemical and structural data showing that Pol I handles Ap sites in a unique manner. Contrary to CPD lesions, Ap sites can template preferential addition of adenine opposite the lesion, following the A-rule. Nevertheless, in contrast to Pol II, Pol I stalls firmly after nucleotide incorporation opposite the Ap site with minimal bypass. Moreover, our cryo-EM structures, at resolutions ranging from 2.8 to 3.5 Å, capture cleft opening and A12-Ct access in the enzyme funnel next to the active center, provide the structural basis for adenosine addition opposite the Ap site, present atomic evidence for subsequent stalling, and uncover the presence of an E site in Pol I. Altogether, our results shed light on the mechanism of Ap site recognition and handling by Pol I.

## Results

### Pol I stalls at abasic sites and cleaves slowly-inserted purines

To explore the effect of Ap sites on Pol I elongation, we performed in vitro transcription tests using a nucleic acid scaffold that contained an Ap site analog (Fig. 1, central inset) in the template DNA strand. The scaffold includes an 8-mer RNA whose 3’-end hybridizes with the template strand one nucleotide before the Ap site (Fig. 1A). Upon addition of NTPs to the mix, Pol I slowly incorporates one nucleotide opposite the lesion but further RNA extension is strongly disfavored. While nucleotide incorporation opposite the Ap site is also slow in the case of Pol II, this enzyme elongates the RNA molecule more efficiently than Pol I, in agreement with previous reports^9,35^. Besides, in the case of Pol I, we observed that its intrinsic RNA cleavage activity partially degrades the RNA molecule and that this effect is reduced in the presence of NTPs.

**Figure 1.**
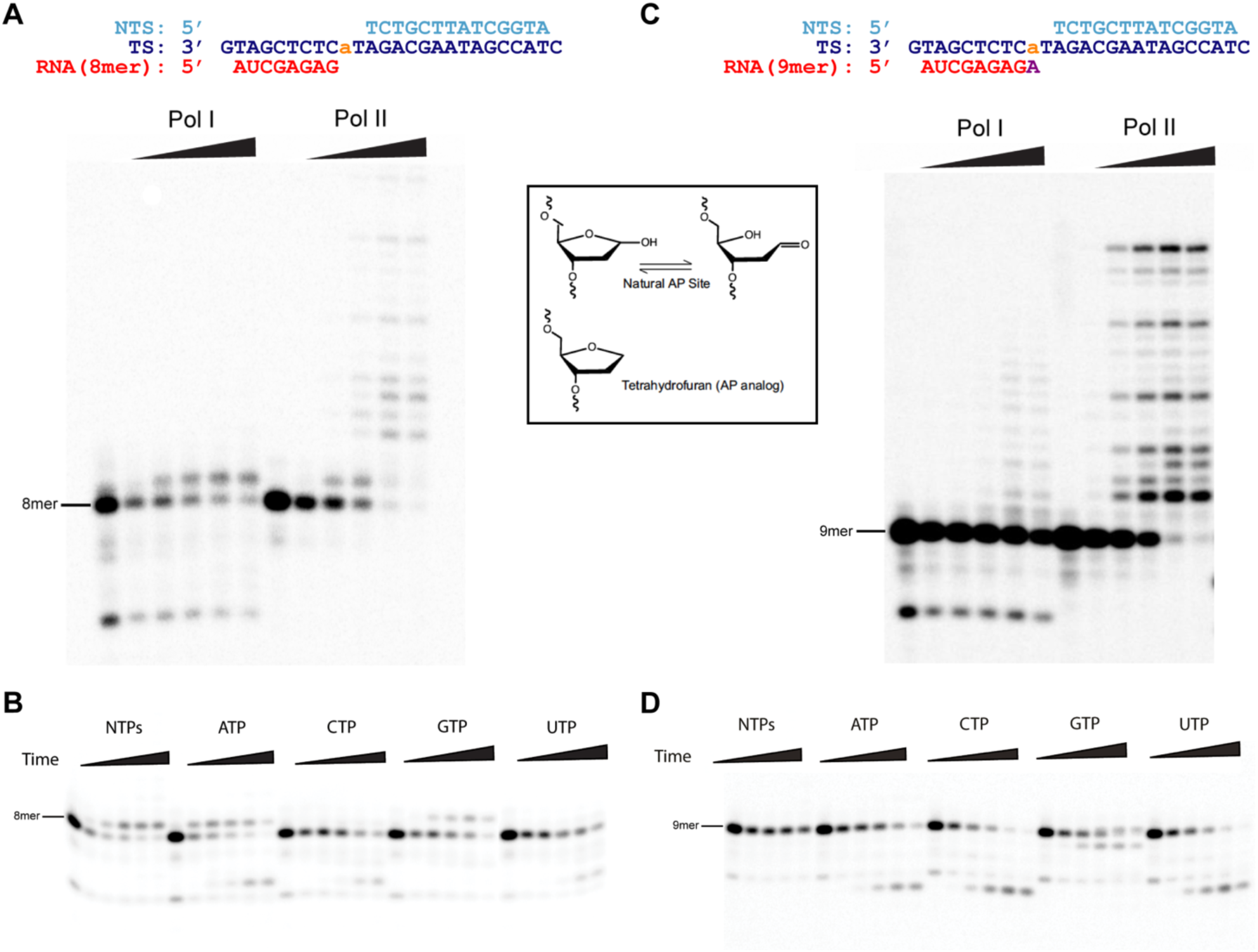
Ap site has distinct effects on Pol I and Pol II elongation. The central inset shows a natural Ap site equilibrium between the closed sugar hemiacetal and the open aldehyde and, below, the synthetic analog tetrahydrofuran used in this study. (A) In vitro RNA extension assay on a scaffold containing an Ap site at *i+1*. Each lane corresponds to incubation times of 0, 0.25, 1, 4, 15 and 60 min. (B) RNA extension assay with the same scaffold as in panel A but using individual NTPs. (C) RNA extension assay on a scaffold containing an Ap site at *i–1*. (D) RNA extension assay with the same scaffold as in panel C but using individual NTPs. Each lane corresponds to incubation times of 0, 1, 4, 15 and 60 min.

We then investigated the effect of Ap sites on nucleotide selectivity and found that Pol I follows the A-rule for nucleotide addition opposite the Ap site (Fig. 1B). Interestingly, guanosine can also be incorporated though at a slower rate, while pyrimidine incorporation is strongly disfavored. Our measured dNTP selectivity of Pol I opposite an Ap site is 1A:0.5G:0.1T:0.1U. Notably, we detected an RNA cleavage product upon prolonged incubation with individual NTPs, with the exception of GTP or when all NTPs are present. This prompted us to perform in vitro transcription tests using a nucleic acid scaffold containing adenosine opposite an Ap site, which is followed by thymine as DNA templating nucleotide (Fig. 1C). Lack of base pairing at the RNA 3’-end likely involves an anomalous configuration of the RNA/DNA hybrid. In these conditions, Pol II is able to bypass the lesion at slow rate, as reported^9^. In contrast, Ap site bypass is negligeable in the case of Pol I, while RNA cleavage is relevant. Incubation with cognate ATP or non-cognate pyrimidines led to formation of 5-mer RNA products, although accumulation of the cleavage product is slower in the case of ATP (Fig. 1D). Incubation with GTP led to formation of a mix between 8-mer and 9-mer RNAs, due to the fact that GTP is the cognate for the DNA nucleotide upstream of the Ap site, suggesting a balance between RNA elongation and cleavage at this position. Overall, our results show that Pol I slowly inserts a purine opposite Ap sites, with preference for adenine, but its intrinsic RNA cleavage activity minimizes lesion bypass.

### Structure of Pol I paused at an abasic site

To uncover the structural basis of Pol I stalling at Ap sites, we prepared Pol I elongation complexes using a mismatched transcription bubble containing an Ap site at the DNA templating position (*i+1*) (Fig. 2A). A non-hydrolysable ATP analog (AMPCPP) was included in the mixture to mimic the insertion step of nucleotide addition opposite the Ap site. We applied cryo-EM to obtain an overall map at 2.8 Å resolution (Map-I), with regions attaining 2.6 Å resolution at the Pol I core including the active center (Fig. S1-S3). The structure, hereafter termed EC-I (Fig. 2B), showed that Pol I adopts a post-translocated configuration with the Ap site at the *i+1* position (Fig. 2C). Nevertheless, the configuration of the DNA backbone around the Ap site differs from that observed in a post-translocated Pol I EC with undamaged DNA^36^ (Fig. 2D). The Ap site ribose is rotated by ∼60° towards the bridge helix and forms a hydrogen bond (H-bond) with the main chain of the bridge helix residue S1014, associated to a minor distortion of the bridge helix in the vicinity of this residue (Fig. 2E). Moreover, in contrast to Pol I EC with undamaged DNA, R1070 in A135 loop and K462 in switch loop 2 of A190 contact the DNA mainchain. Besides, hybrid base pairs at positions *i–1* and *i–2* are tilted by 30° towards the bridge helix, likely due to base absence at the Ap site (Fig. 2D-E). Superposition with the structure of Pol I stalled at a CPD lesion shows that the template DNA strand lies backwards in the CPD lesion structure, which affects the configuration of the DNA/RNA hydrid (Fig. 2F). Importantly, while the Ap site surpasses the bridge helix barrier and occupies the templating position, the CPD lesion lies behind the bridge helix. This explains that RNA extension by Pol I stops before the CPD lesion^34^, while Ap sites induce enzyme stalling after slow nucleotide addition opposite the lesion (Fig. 1).

**Figure 2.**
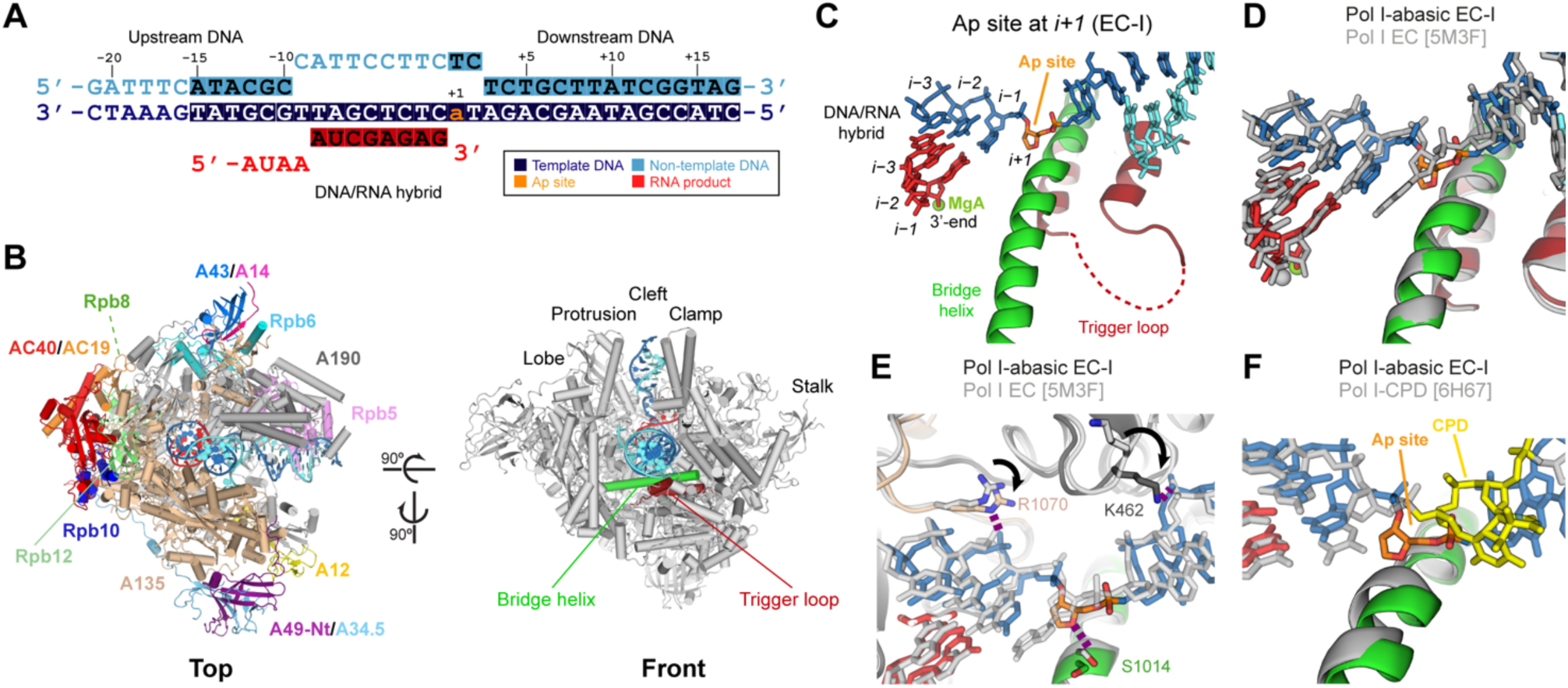
Structure of Pol I paused by an Ap site at *i+1*. (A) Schematic diagram of the nucleic acid scaffold with an Ap site at *i+1*. Filled squares denote nucleotides visible in the cryo-EM map that were modeled. (B) Two views of Pol I paused at an Ap site (EC-I) indicating the different subunits and structural elements in the enzyme. (C) Close-up view of the active site of the EC-I structure. (D, E) Two views of the comparison between EC-I (colors) and Pol I with undamaged DNA in a post-translocated state (PDB code 5M3F, grey). H-bonds are shown as purple dotted lines. Arrows in panel E indicate residue changes between ECs with undamaged and damaged DNA. (F) Superposition of EC-I (colors) with Pol I stalled at a CPD lesion (PDB code 6H67, grey and CPD lesion in yellow).

### The A12 C-terminal Zn ribbon inserts in the funnel to catalyze RNA cleavage

To shed light on conformational changes of Pol I paused at Ap sites, we conducted an overall 3D classification of the cryo-EM particles. One of the classes showed additional density for A12-Ct bound to the enzyme funnel. Focused 3D classification using a mask covering the funnel yielded an improved map with an overall resolution of 3.3 Å, hereafter Map-II (Fig. S1-S3). The complete A12 subunit including both ZnR (residues 1-43 and 80-125) and the connecting linker could be modelled to obtain EC-II (Fig. 2A-B), in spite of poor density for the catalytic loop harboring acidic residues D105 and E106. Insertion of A12-Ct in the funnel has been observed in Pol I devoid of nucleic acids^19,20,36^ or in a pre-initiation complex lacking RNA^37^ (Fig. 3C). Accommodation of A12-Ct within the funnel involves partial opening of the DNA-binding cleft by 30 Å, associated to unfolding of the central region in the bridge helix (Fig. 3D-E). A similar configuration has been reported for Pol II complexed to TFIIS^13^ and for Pol III in complex with the integrase of the Ty1 retrotransposon^38^. Additionally, residue Y717 in A135 moves away from the A12 acidic loop to avoid clashes (Fig. 3B). The equivalent residue in Pol II, Y769 in subunit Rpb2, was proposed to limit the extent of backtracking^13^. The overall configuration of the hybrid is similar to EC-I, while the Ap site and downstream DNA accompany the clamp as the cleft opens (Fig. 3F). Therefore, the EC-II structure suggests that an Ap site at the templating i+1 position induces an RNA cleavage-prone configuration in Pol I and supports residual RNA cleavage products in our RNA extension assays (Fig. 1B-D). The EC-II structure may also represent a post-cleavage configuration in which A12-Ct has not abandoned the funnel.

**Figure 3.**
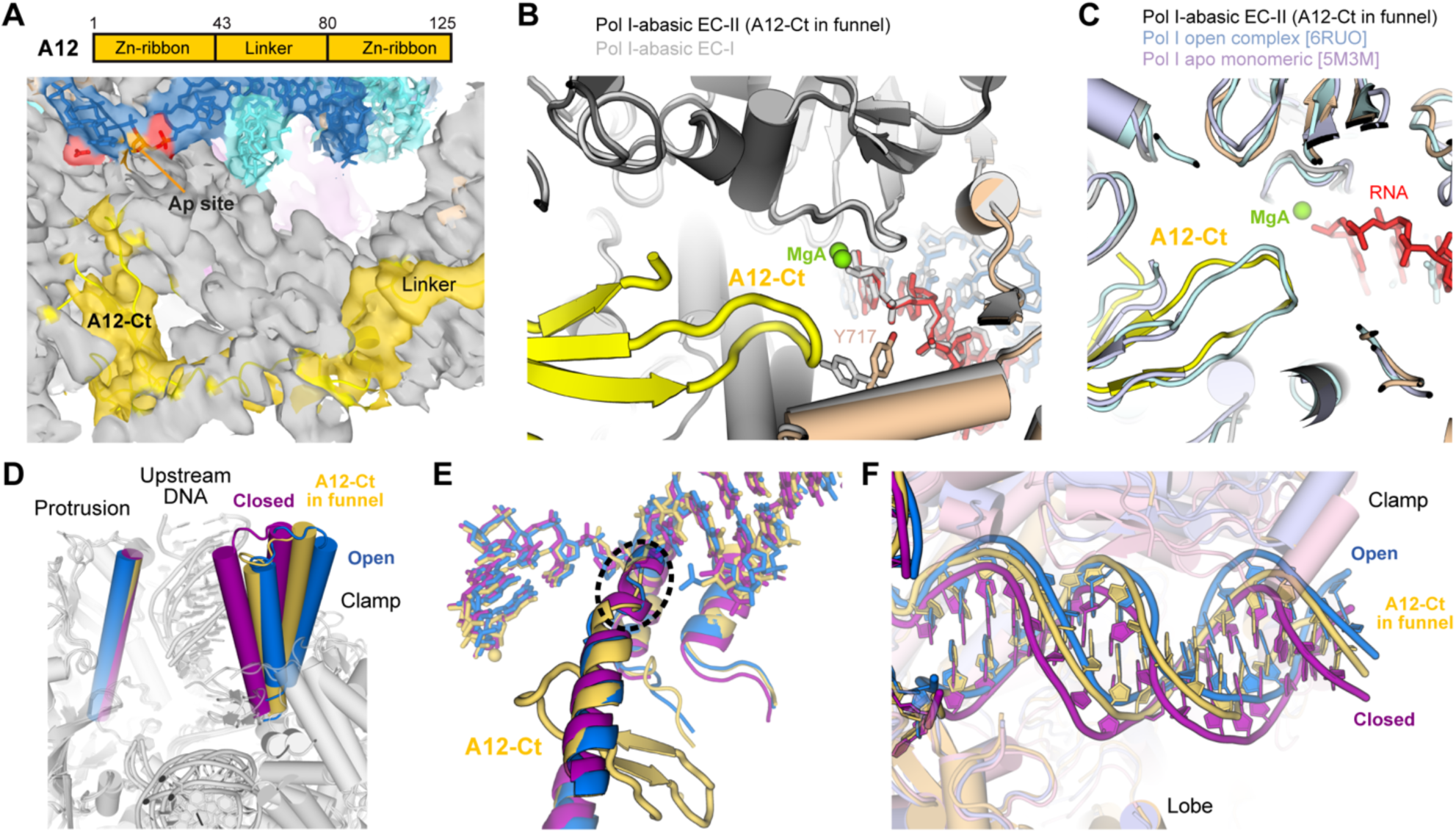
Paused Pol I by Ap site at *i+1* with A12-Ct in the funnel. (A) Bar diagram of subunit A12 and EC-II cryo-EM map showing density for A12-linker and A12-Ct. The template and non-template strands are in blue and cyan, respectively. (B) Superposition of EC-II (colors) with EC-I (grey). The A12-Ct catalytic loop (residues 103-108) in EC-II was modeled in poor density, as seen in panel C. (C) Superposition of EC-II (colors) with a Pol I open complex where A12-Ct occupies the funnel (PDB code 6RUO, aquamarine) and free monomeric Pol I with A12-Ct in the funnel (PDB code 5M3M, violet). (D-F) Superposition of EC-II (yellow) with EC-Ia (purple) and EC-Ib (blue) in the front view at the cleft entrance (panel D), around the active center (panel E), and in the top view around downstream DNA (panel F). The dotted circle in panel F indicates a bridge helix region that partially unfolds upon cleft opening.

Two additional classes produced maps that are equivalent to Map-I but show differences in the DNA-binding cleft configuration (Fig. S1-S3). Map-Ia presents a closed cleft while Map-Ib exhibits an open cleft, and both exhibit a resolution of 3.5 Å (Movie S1). Comparison of the resulting structures shows that cleft opening depends on pivoting along the downstream DNA axis (Fig. 3D), with minimal aperture of 28 Å in EC-Ia and maximal aperture of 32 Å in EC-Ib. This is consistent with previous observations^20,39^. The Ap site in the closed state is retracted respect to EC-I, while it is further inserted in the active site in the open state (Fig. 3F), while downstream DNA moves with the clamp during cleft opening (Fig. 3E). This indicates that we have captured different intermediate states of Ap site translocation into the Pol I active center prior to NTP insertion. In the open state, densities for both upstream and downstream DNA are poor, while domains several domains appear flexible, including half of the clamp and part of the jaw in A190, most of the stalk, and the C-terminal region of Rpb5. Therefore, we interpret this open configuration as a Pol I paused state that may be prone to DNA dissociation.

### Structures of Pol I lacking lobe-associated subunits

Two additional classes lacked density for subunits binding next to the A135 lobe. Map-III, showing a resolution of 3.2 Å, corresponds to Pol I* paused at an Ap site as it lacks density for the A49/A34 heterodimer (Fig. S1-S3). The nucleic acid configuration in the resulting EC-I* structure is virtually identical to that observed in EC-I (Fig. 4A). In the absence of A49/A34.5, A12-Ct binds external domains 1 and 2 (ED1 and ED2), a region in the A135 lobe that is hindered in the presence of A49/A34.5 (Fig. 4B). This A12-Ct location is equivalent to that observed for the Pol I* EC with undamaged DNA^33^, which is overall similar to EC-I* except that downstream DNA lies backwards in the latter (Fig. 4B). Comparison with the EC-II structure shows that A12-Ct undergoes a movement of ∼60 Å and a rotation of ∼90°, using the central mini-helix (residues 59-65) in the A12-linker as a hinge.

**Figure 4.**
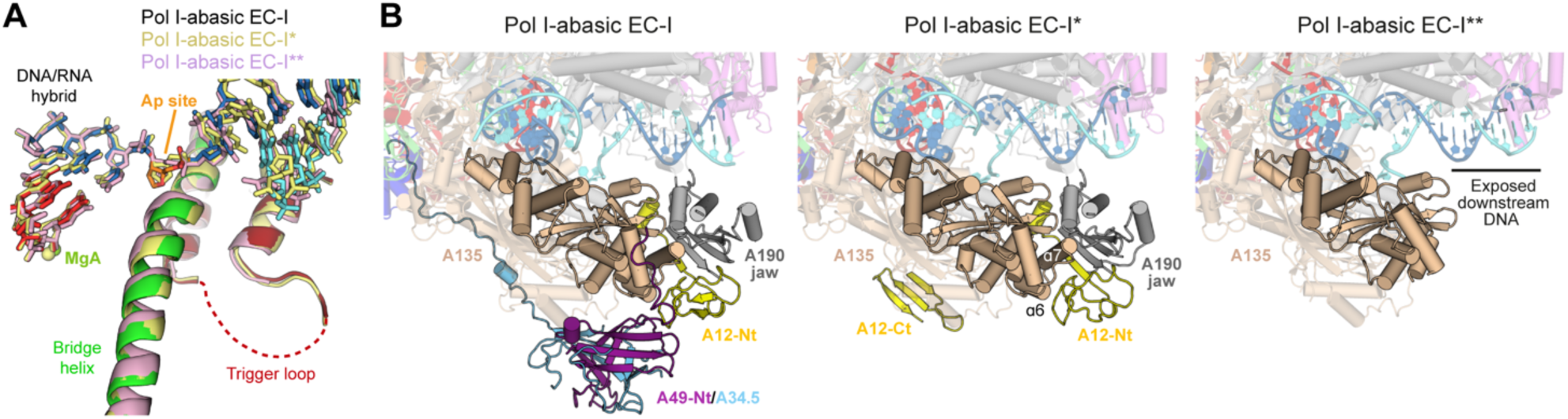
Structures of Pol I paused by an Ap site at *i+1* lacking lobe-associated subunits. (A) Close-up view around the active center of the superposition of EC-I with EC-I* and EC-I**. The bridge helix and trigger loop in EC-I are in green and red. (B) Overall top view of EC-I, EC-I* and EC-I**.

A final class arising from overall 3D classification produced Map-IV, which presents an overall resolution 3.5 Å and lacks subunits A49, A34 and A12 (Fig. S1-S3). Unexpectedly, the resulting EC-I** structure not only lacks density for these three subunits, but also for the entire A190 jaw domain (residues 1241-1540) and helices α6-α7 in the A135 lobe (residues 281-324) (Fig. 4B). This indicates that the N-terminal ZnR in A12 (A12-Nt) plays a stabilizing role of the A190 jaw and part of the lobe, which become flexible when A12-Nt is absent. As a result, about two thirds of downstream DNA become exposed to the solvent, which may allow access of DNA-binding proteins into the Pol I cleft (Fig. 4B). Surprisingly, the conformation of the enzyme is similar to that observed in EC-I or EC-I*, but the cleft is slightly more closed in EC-I**. This suggests that the jaw is able to move independently using the cleft-jaw link as a hinge. Moreover, the nucleic acid scaffold also adopts an equivalent configuration, suggesting that this Pol I form may be able to transcribe in vivo, consistent with cell viability of a ΔA12 strain where the Pol I enzyme lacks the three lobe-associated subunits^40,41^.

### Purine addition opposite the abasic site occurs via base stacking

Focused 3D classification with a spherical mask around the active center identified two classes with density for AMPCPP. The refined Map-Va and Map-Vb attained resolutions of 3.0 and 3.3 Å, respectively (Fig. S1-S3). The former exhibits clear density next to the RNA 3’-end corresponding to the adenine and phosphate moieties of AMPCPP, while the ribose appears flexible (Fig. S3). In the resulting EC-IIIa structure, the adenine moiety of AMPCPP is sandwiched between the base at the RNA 3’-end and residue T1013 within the bridge helix (Fig. 5A). P593 in subunit A190, a residue that is strictly conserved among nuclear RNA polymerases (Fig. S4), lies at hydrophobic distance from adenine, likely explaining the observed preference versus guanine (Fig. 1B). R714 and R957 in subunit A135, both strictly conserved among nuclear RNA polymerases (Fig. S5), lie at H-bond distance from the γ-phosphate. The active center configuration is equivalent to that obtained from all particles, except that the Ap site moves away from the canonical templating position (Fig. 5B). Comparison with Pol I EC with undamaged DNA containing GMPCPP^33^ shows that AMPCPP in EC-III occupies the canonical NTP addition site, i.e. the A-site (Fig. 5C). However, the bridge helix approaches the incoming nucleotide to position the T1013 sidechain for optimal purine sandwiching, while the trigger loop is retracted and can be fully modelled (Fig. 5B). Overall, this unique configuration of the active site is likely derived from lack of templating base at the Ap site and establishes the molecular basis of the A-rule for Pol I paused at Ap sites.

**Figure 5.**
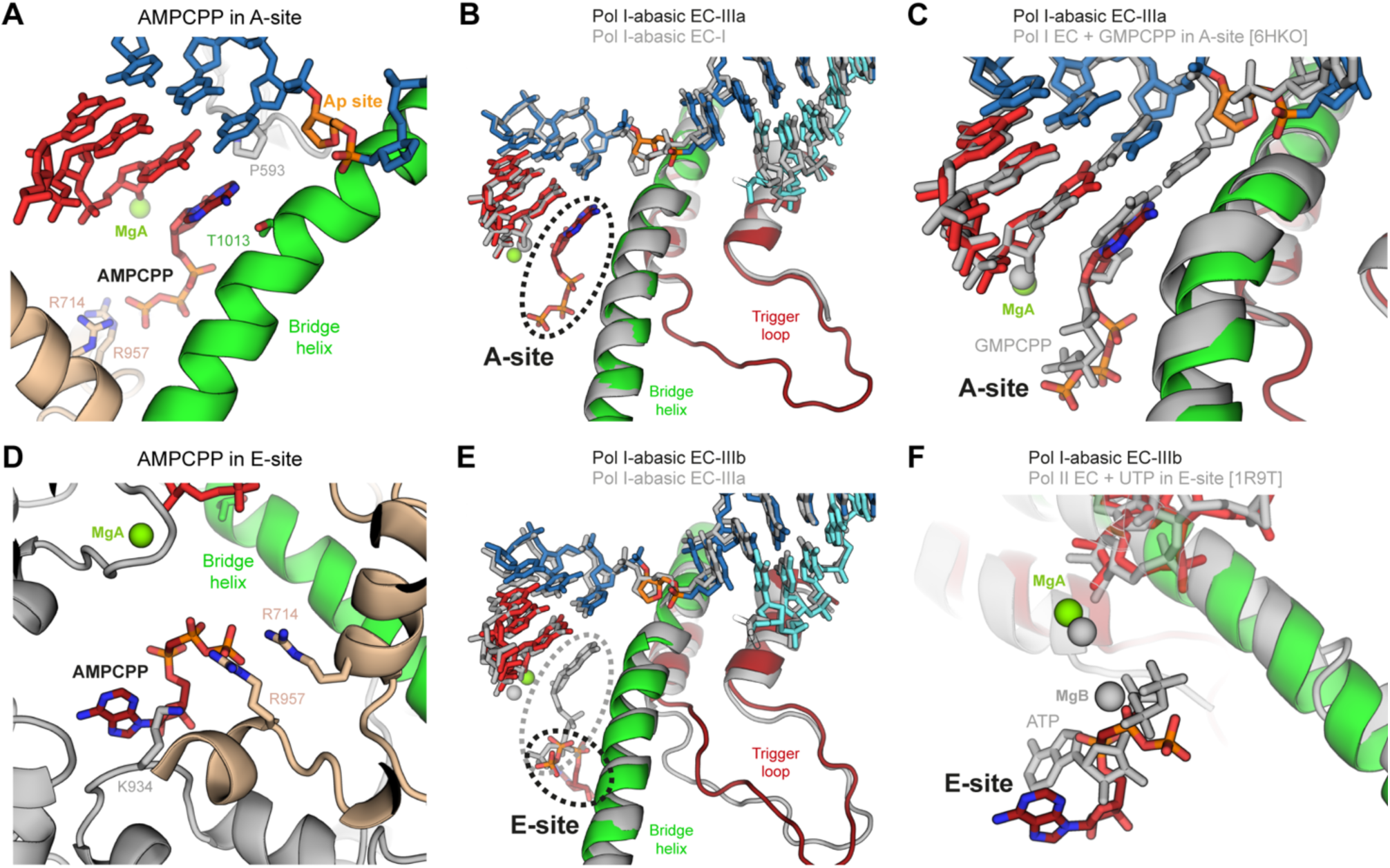
Structures of Pol I paused by an Ap site at *i+1* in the presence of AMPCPP. (A) Close-up view around AMPCPP in the A-site. (B) Superposition of EC-IIIa (colors) with EC-I (grey). (C) Superposition of EC-IIIa (colors) with Pol I with undamaged DNA in a pre-insertion state in the presence of GMPCPP in the A-site (PDB code 6HKO, grey). (D) Close-up view around AMPCPP in the E-site. The view is rotated ∼120° along the bridge helix respect to that in panel A. (G) Superposition of EC-IIIb (colors) with EC-IIIa (grey). (H) Superposition of EC-IIIb (colors) with Pol II with ATP in the E-site (PDB code 1R9T, grey). View is as in panel D.

Map-Vb exhibits density for AMPCPP in an alternative location beneath the active site (Fig. S3). In the resulting EC-IIIb structure, the adenine moiety of AMPCPP lies next to K934 in A190, while the γ-phosphate forms H-bonds with R714 and R957 in A135 (Fig. 5D-E). These three residues are strictly conserved among nuclear RNA polymerases (Fig. S4-S5). We assign this location to the NTP entry site, i.e. the E-site, previously described in a Pol II EC structure in complex with a non-cognate NTP^15^ (Fig. 5F). As in Pol II, the β- and γ-phosphates binding sites overlap in the A- and E-sites, implying that they cannot be occupied simultaneously (Fig. 5E). However, in spite of similarities between the two enzymes, the Pol I E-site appears to bind the NTP base more tightly, which may be due to absence of MgB in our structure. In our structure with AMPCPP in the E-site, the bridge helix is partly unfolded next to the Ap site, while the trigger loop is retracted and can be fully modelled (Fig. 5G). Additionally, DNA around the Ap site appears flexible due to weak density in this region of the map (Fig. S3).

### Structure of Pol I stalled after addition opposite an abasic site

Finally, we studied the cryo-EM structure of Pol I stalled after addition of adenine opposite the Ap site, using a nucleic acid scaffold where the Ap:adenine unpaired duo is placed at the *i–1* position (Fig. 6A). Focused 3D classification with a mask covering the DNA-binding cleft identified a population where the Ap:adenine unpaired duo is clearly defined (Fig. S1-S3). To our surprise, the refined 2.8 Å Map-VI corresponds to Pol I*, i.e. lacking A49/A34.5 and with A12-Ct bound on the lobe. In the derived EC-IV structure (Fig. 6B) nucleic acids adopt a configuration that is compatible with an intermediate of translocation (Fig. 6C). Notably, the DNA base at position *i–2* is tilted by ∼30° so that it forms H-bonds with RNA bases at positions *i–2* and *i–1*, the latter being otherwise unpaired due to lack of base in the opposite Ap site (Fig. 6D-F). Equivalently, the DNA base at position *i–3* is tilted so that it forms H-bonds with RNA bases at positions *i–3* and *i–2*. The position of the Ap site is clearly defined, slightly advanced respect to EC-I and EC-IIIa (Fig. 6D-E) but far from the post-translocated state (Fig. 6F). Downstream DNA also presents an intermediate state of translocation and the *i+1* nucleotide lies behind the bridge helix, such that it cannot template nucleotide addition (Fig. 6C). RNA nucleotides are closer to the post-translocated state but lie 1.6 Å behind the canonical position, thus partly occluding the A-site. While the central region of the bridge helix is fully folded, it lies closer to the trigger loop respect to the post-translocated EC-I structure, leading to slight opening of the cleft. Overall, this provokes a less tighter interaction of the template DNA with Pol I and may be related to lack of subunits A49 and A34.5. In conclusion, EC-IV presents an intermediate translocation state that is disfavored for subsequent RNA elongation. Nevertheless, this configuration is different from the bacterial RNA polymerase swiveled state^42^ or the mammalian Pol II tilted state^43^. In particular, the central region of the bridge helix lies ∼3 Å closer to the trigger loop, while DNA bases in *i–2* and *i–3* occupy the space of *i–1* and *i–2* bases in other paused states (Fig. 6G). This suggests that Pol I stalls at Ap sites through a unique mechanism.

**Figure 6.**
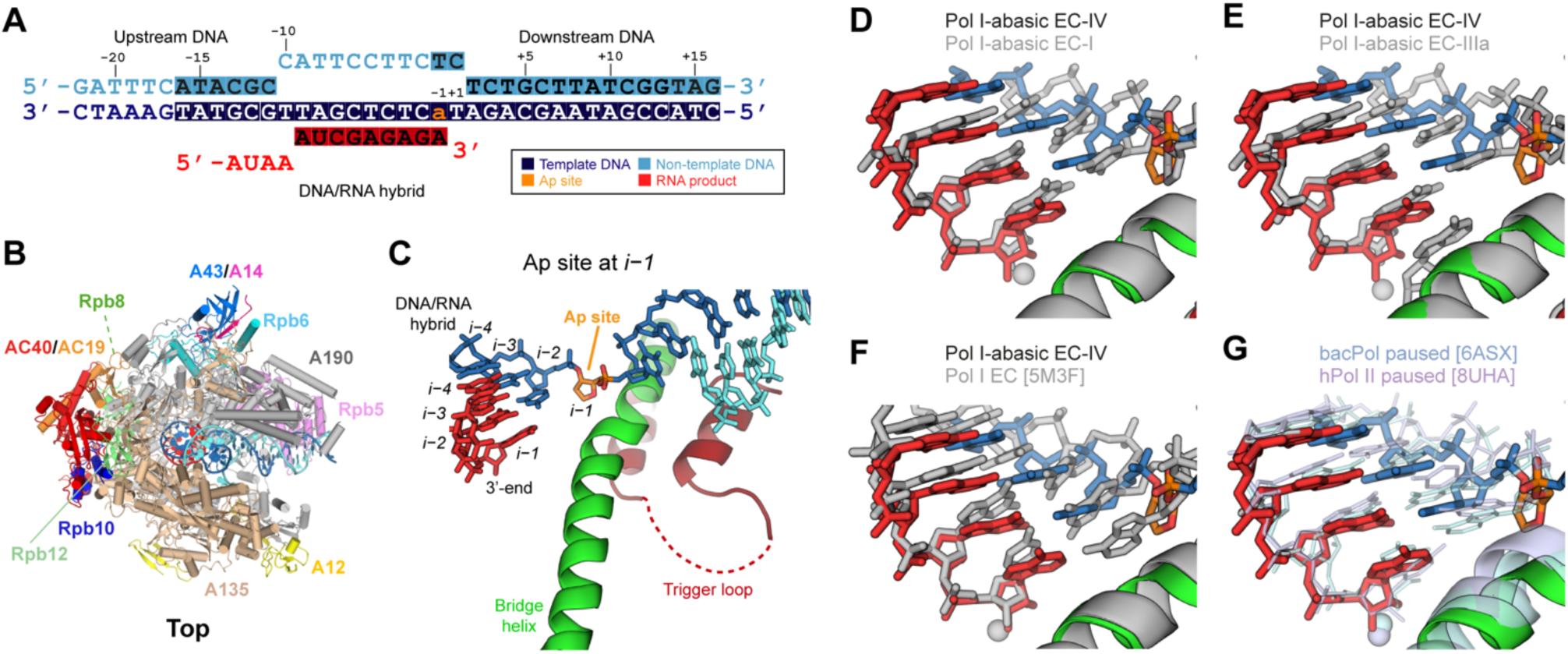
Structure of Pol I stalled by an Ap site at *i–1*. (A) Schematic diagram of the nucleic acid scaffold with an Ap site at *i–1*. Filled squares denote nucleotides visible in the cryo-EM map that were modeled. (B) Overall view of Pol I stalled at an Ap site (EC-IV) indicating the different subunits. (C) Close-up view of the active site in the EC-IV structure. (D-G) Close-up views comparing the position of nucleotides in the DNA/RNA hybrid between EC-IV (colors) and EC-I (grey, panel D), EC-IIIa (grey, panel E), Pol I EC with undamaged DNA (PDB code 5M3F, grey, panel F), paused bacterial RNA polymerase (PDB code 6ASX, aquamarine; panel G), or paused human Pol II (PDB code 8UHA, violet; panel G).

## Discussion

DNA lesions on the template strand challenge transcription elongation by slowing or blocking the advance of RNA polymerases. In this work, we reveal the detailed mechanism of Pol I stalling at Ap sites, which are the most common DNA lesions in living cells, by combining biochemical analysis with cryo-EM structures of several conformational states in the process. Our results allow us to propose a pathway for Ap site-induced pausing and stalling that consists of several steps (Fig. 7). Contrary to CPD lesions, which induce firm Pol I stalling before nucleotide addition opposite the lesion^34^, Ap sites allow for untemplated purine addition opposite the lesion at slow rate, followed by Pol I stalling with minimal bypass after prolonged incubations. Importantly, similar to CPD lesions, the intrinsic RNA cleavage activity residing in A12-Ct dramatically reduces Ap site bypass by Pol I.

**Figure 7.**
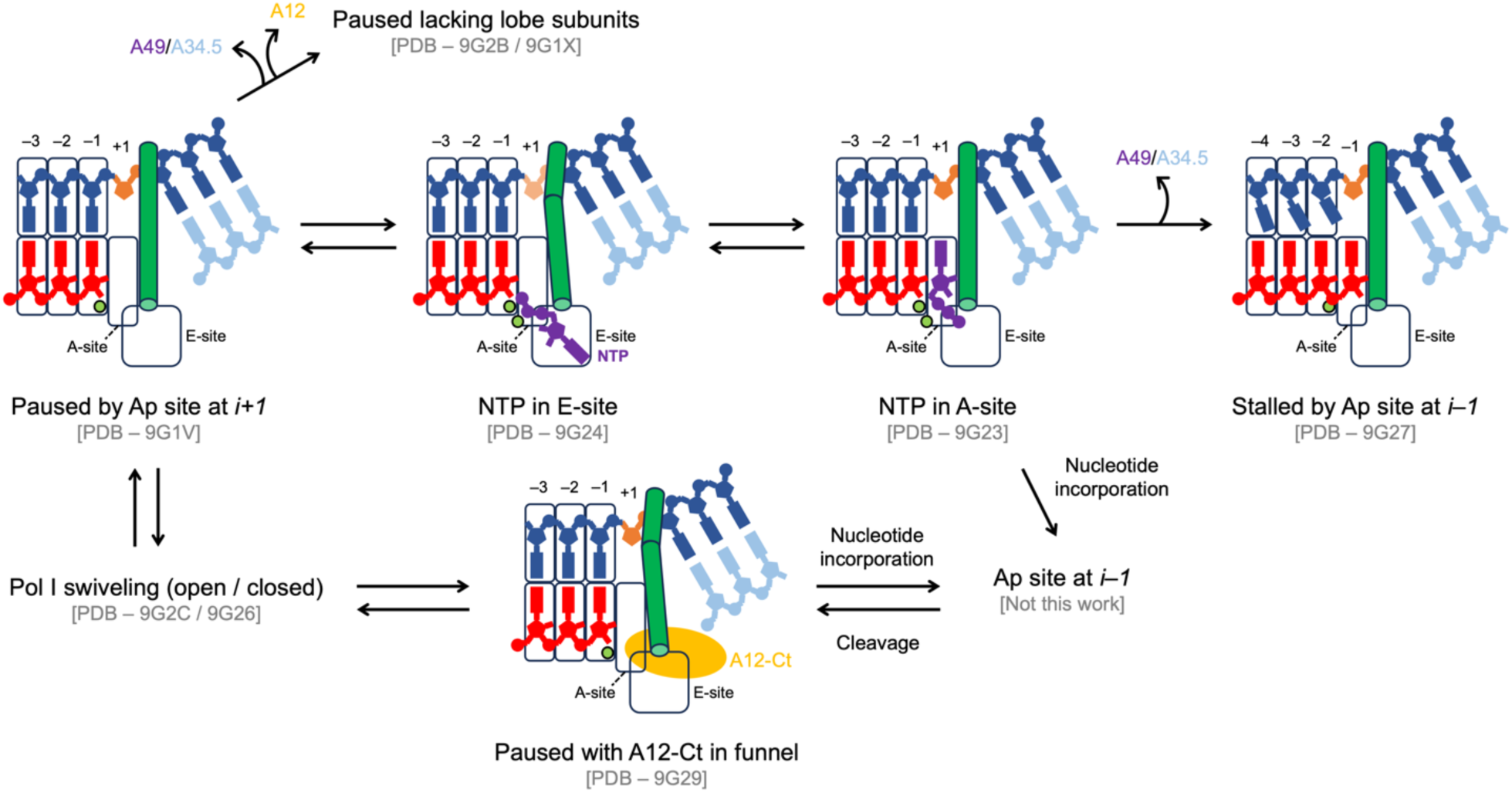
Mechanistic model of Pol I pausing and stalling at Ap sites. The DNA template and non-template strands are in blue and cyan, RNA is in red, the Ap site is in orange, the incoming nucleotide is in purple, A12 is in yellow, the bridge helix is in dark green and magnesium ions are in light green. Pol I is initially stalled as the Ap site occupies the templating position (*i+1*), which leads to cleft swiveling. This may allow access of A12-Ct into the funnel for subsequent RNA cleavage or lead to Pol I dissociation from DNA in the open cleft state. Pol I stalled at Ap site allows NTP entry into the E-site, which enables access into the A-site. In the A-site, purines are stabilized by sandwiching between the RNA 3’-end and the bridge helix, with preference for ATP. Phosphodiester bond formation likely leads to an altered hybrid configuration that induces RNA cleavage to minimize lesion bypass. Alternatively A49/A34 are lost, which hampers RNA cleavage but induces an intermediate of translocation that compromises nucleotide addition, thus stalling Pol I.

### Pol I swiveling at templating Ap sites

Our structures of Pol I with an Ap site at *i+1* show that nucleic acids in the cleft adopt a post-translocated configuration where the lesion occupies the templating position. Nevertheless, the absence of a templating base induces changes in the hybrid, the bridge helix and switch loop, which likely make this post-translocated state suboptimal for addition. Consistently, two conformations of the cleft aperture are observed, open and closed (Movie S1), suggesting that the lack of a templating base is sensed by Pol I. In the open state, the downstream and upstream regions of DNA are loosely bound to the enzyme, which may lead to Pol I fall off from DNA at templating Ap sites. In agreement, the Pol I elongation complex presents low stability as compared to other RNA polymerases^44^.

Alternatively, cleft opening may be necessary to allow access of A12-Ct into the funnel. This is supported by our structure of Pol I with A12-Ct in the funnel, which exhibits partial cleft opening, as previously observed for an initiation open complex in the absence of RNA^37^. Our structure indicates that lack of a templating base may induce a configuration that is prone to RNA cleavage, although our biochemical assays reveal that RNA cleavage mainly occurs after nucleotide addition opposite the Ap site. Importantly, location of A12-Ct in the funnel leaves a channel for NTP entrance into the active site (Fig. 3A). Moreover, flexibility of the A12-Ct acidic loop in our structure likely allows NTP binding to the E- or the A-site for nucleotide addition.

### Purine addition occurs through base sandwiching

The structures of Pol I paused at Ap sites complexed to AMPCPP uncover the E- and A-sites for nucleotide incorporation opposite abasic lesions. The NTP in the E-site binds mainly through its terminal diphosphate moiety to two arginine residues in subunit A135 that are conserved among nuclear RNA polymerases. Interestingly, the NTP base forms stabilizing interactions with a lysine in A190 that is conserved in all RNA polymerases. A pyrimidine is likely to establish equivalent weak interactions with this Pol I region. A second Mg ion, as seen in Pol II^9,15^, is likely to further stabilize the interaction on the phosphate moiety and favor rotation into the A-site. The existence of an E-site in Pol I may be related to transcriptional speed, as shown for Pol II^45^. However, the Pol I and Pol II E-sites differ in the position of the NTP base, with stronger binding in the case of Pol I. This may enhance NTP residence in the funnel, consistent with faster elongation rates of Pol I respect to Pol II^44,46^.

In the absence of a DNA templating base, NTP in the A-site is able to occupy the canonical position, thus enabling addition opposite the Ap site, as shown in our RNA extension assays. This is possible due to NTP base sandwiching by the bridge helix and the base at the 3’-end of the RNA, which explains the preference for purines as their stacking capacity is higher than for pyrimidines. While our results were obtained with a nucleic acid scaffold harboring a purine base at the RNA 3’-end, an equivalent effect is expected if a pyrimidine base is present at the RNA 3’-end. The minor preference of adenine versus guanine is likely due a conserved proline next to the active center.

### Stalling after purine addition is incompatible with RNA extension

According to our results, NTP addition opposite Ap sites leads to Pol I translocation stalling, which affects both the template DNA strand and the RNA. On the RNA side, incomplete translocation leads to partial occlusion of the A-site. On the DNA side, the *i+1* nucleotide lies behind the bridge helix, hampering its templating role. This is due to stabilization of the unpaired base at the RNA 3’-end by the DNA base in the hybrid that is adjacent to the Ap site. Importantly, the intermediate translocation state observed in our structure differs from described translocation intermediates^42,43^. In this unique paused state, which we term retracted pause, nucleotide addition is compromised, in agreement with our RNA extension assays (Fig. 1). Moreover, these assays suggest that the retracted pause induces Pol I backtracking and RNA cleavage, leading to persistent stalling after nucleotide addition opposite Ap sites, with minimal bypass upon long incubations.

### Role of lobe-associated subunits A12 and A49/34.5

Lobe-associated subunits A49, A34.5 and A12 provide distinctive functions to Pol I^47^ in spite of being dispensable for yeast growth^40,41,48^. We find that the A49/A34.5 heterodimer can dissociate from Pol I when an Ap site in the template DNA strand is positioned at the *i+1* or *i–1* site. This heterodimer was proposed to reversibly bind the Pol I core with a putative in vivo role in initiation and elongation^33^. Our data support this hypothesis but further suggest a role in DNA lesions detection or, more generally, in transcriptional pausing. While we show that backtracking and RNA cleavage are induced at Ap sites, Pol I is persistently stalled after NTP incorporation opposite the lesion due to a non-productive translocation intermediate. A paused Pol I configuration may induce dissociation of A49/A34.5, which would otherwise travel with Pol I during rRNA synthesis. This is consistent with ChIP data showing that the heterodimer associates with the entire pre-rRNA gene^49,50^.

A49/A34.5 dissociation has two major effects on the Pol I EC. On one hand, the A12-Ct binding site at Pol I external domains becomes exposed, leading to A12-Ct fixation on the lobe. This is consistent with reduced RNA cleavage activity respect to the complete enzyme^30^. On the other hand, the binding site for the C-terminal domain of A49 next to upstream DNA becomes available. In Pol II, processivity factor Spt4/5 has been shown to bind in the same region^51^ and Spt5 was shown to bind Pol I^52^. These two effects could contribute to increased lesion bypass. Interestingly, as opposed to Pol I stalled at CPD lesions^34^, no density for A49-Ct is observed in particles containing A49/A34.5. This suggests that bypass of Ap sites may be favored respect to CPD lesions.

Unexpectedly, we found particles that lack all three lobe-associated subunits when an Ap site occupies the templating position. These structure shows a highly-exposed downstream DNA due to flexibility of the A190 jaw domain and a pair of helices in the lobe. This may enable access to factors that bind downstream DNA for repair. Nevertheless, the structure is compatible with nucleotide addition, consistent with studies showing that the presence of A12 destabilizes the Pol I elongation complex^53^. Moreover, biochemical analysis of Pol I lacking the three lobe-associated subunits showed that this form of the enzyme is deficient in NTP selection and induces errors in the transcript^30^. Therefore, loss of lobe-associated subunits likely favors Ap site bypass.

### Different stalling mechanisms in Pol I and Pol II

Pol I stalling at Ap sites follows a two-step mechanism (Fig. 7). In the first step, Pol I is able to slowly add a nucleotide opposite the lesion, with preference for adenosine, which is equivalent to Pol II^9^. In the second step, Pol I stalls firmly due to impaired nucleotide addition and fast RNA cleavage, leading to minimal bypass. In contrast, Pol II has a strong tendency to bypass Ap sites after nucleotide addition opposite the lesion. On one hand, this is due to the fact that Pol II is able to fully translocate after nucleotide addition opposite the lesion, while translocation is unfavored in Pol I (Fig. 6). On the other hand, the intrinsic Pol I cleavage activity acts as failsafe mechanism to minimize bypass, while Pol II requires TFIIS binding and efficiency is reduced. Our results suggest that Ap site bypass by Pol I is more deleterious for the cell than bypass by Pol II, as described for CPD lesions^34^.

While the main pathway to repair Ap sites is BER, transcription-coupled nucleotide excision repair (TC-NER) has been reported to operate for Ap site removal from the template DNA strand in yeast^54^. Moreover, NER was shown to repair Ap sites in genes that are highly transcribed by Pol II^35,55,56^. Given rDNA is highly transcribed, it is plausible that a similar mechanism operates for Pol I, as this pathway was shown to operate in this transcription system^57^. Our structure of stalled Pol I lacking the three lobe-associated subunits provides accessibility to downstream DNA, that may operate in such mechanism.

## Methods

### Protein purification

Purification of yeast Pol I was performed following described procedures^58^ with minor modifications. A *Saccharomyces cerevisiae* strain harboring a tandem affinity-purification (TAP) tag at the C-terminus of subunit A190 was grown in a fermenter containing 50 L of YPDA for 20 h at 30 °C to an OD_600_ of 5.5, harvested by centrifugation and stored at -80 °C until use. Subsequent steps were performed at 4 °C. 500 g of cells were resuspended in a buffer containing 250 mM Tris-HCl pH 7.4, 20% glycerol, 250 mM ammonium sulfate, 1 mM EDTA, 10 mM MgCl_2_, 10 µM ZnCl_2_, 10 mM β-mercaptoethanol, supplemented with protease-inhibitors (1 mM phenylmethanesulfonyl fluoride, 2 µg/ml leupeptine, 4 mM benzamidine, 1.4 µg/ml pepstatine A) and DNAse (Roche), and lysed with glass beads using a BeadBeater (Biospec). After centrifugation, the supernatant was incubated with 4 ml of IgG Sepharose (GE Healthcare) for 6 hours, washed with a buffer containing 50 mM Tris-HCl pH 7.4, 5% glycerol, 200 mM NaCl, 1 mM MgCl_2_, 10 µM ZnCl_2_ and 5 mM dithiothreitol (DTT) and incubated overnight with tobacco etch virus (TEV) protease. The TEV eluate was further purified using a Mono Q column (GE Healtcare) and a gradient from 0.2 to 1 M NaCl in buffer 20 mM Tris-HCl pH 7.4, 1 mM MgCl_2_, 10 µM ZnCl_2_ and 5 mM DTT. Pol I-containing fractions were pooled and concentrated to 5 mg/ml, frozen in liquid nitrogen and stored at -80 °C for further use.

### In vitro transcription assays

Non-template DNA (5’-TCTGCTTATCGGTA-3’), template DNA (5’-CTACCGATAAGCAGATaCTCTCGATG-3’) where ‘a’ represents the abasic site, and 8mer RNA (5’-AUCGAGAG-3’) or 9mer RNA (5’-AUCGAGAGA-3’) were purchased from IDT. The transcription elongation assay was performed based on previous reported methods with slight modifications^9^. Briefly, an aliquot of 5’-^32^P labeled RNA was annealed with a 1.5-fold amount of template DNA strand (TS) and 2.0-fold amount of non-template DNA strand (NTS) to form the RNA/DNA mini-scaffold in a buffer composed of 20 mM Tris-HCl (pH 7.5), 150 mM KCl, 5 mM DTT. An aliquot of annealed scaffold was then incubated with a five excess amount of purified Pol I or Pol II on ice for 5 min, followed by incubation at room temperature (23 °C) for 15 min to ensure the formation of Pol I or Pol II elongation complex. The transcription was chased by adding an equal volume of solution containing 150 mM NaCl, 20 mM Tris-HCl (pH 7.5), 5 mM DTT, 10 mM MgCl_2_, and 2 mM NTP. Final reaction concentrations after mixing were 30 nM scaffold, 150 nM Pol I or Pol II, 10 mM MgCl_2_, and 1 mM NTP. Reactions were quenched at various times (0, 0.25, 1, 4, 15, 60 min or 0, 1, 4, 15, 60 min) by addition of one volume of 0.5 M EDTA (pH 8.0). The RNA transcript was analyzed by 12% (wt/vol) denaturing urea/PAGE. The gel was visualized by phosphorimaging and analyzed using ImageLab software (BioRad).

### Preparation of Pol I EC stalled by an abasic site for structural studies

Non-template DNA (5’-GATTTCATACGCCATTCCTTCTCTCTGCTTATCGGTAG-3’), template DNA (5’-CTAAAGTATGCGTTAGCTCTCaTAGACGAATAGCCATC-3’, where ‘a’ corresponds to an Ap site) and RNA-1 (5’-AUAAAUCGAGAG-3’) or RNA-2 (5’-AUAAAUCGAGAGA-3’) from IDT were resuspended in a buffer containing 10 mM Hepes pH 8.0, 150 mM NaCl. The DNA strands were mixed in equimolar amounts, heated to 95 °C and slow-cooled to 4 °C. After, either RNA-1 or RNA-2 was added in equimolar amounts, the mixture heated to 45 °C and slow-cooled to 4 °C. The scaffold was then incubated in equimolar amounts with Pol I overnight at 4 °C in a buffer containing 10 mM Tris pH 7.4, 150 mM NaCl and 5 mM DTT. For the scaffold containing RNA-1, the buffer was supplemented with 5 mM MgCl_2_ and 1 mM AMPCPP. For the scaffold containing RNA-2, 1 mM EDTA was added.

### Cryo-EM sample preparation and data collection

For the sample containing RNA-1, we collected data from two different cryo-EM samples due to strong preferential orientation. For the first dataset, the final Pol I concentration was 0.07 mg/ml and 8 mM CHAPSO was added prior to grid preparation. 3 µl of sample were applied to freshly glow-discharged Quantifoil R2/1, 400 mesh grids coated with a second layer of homemade thin continuous carbon. After 1 min incubation, grids were blotted for 2 s and blot force of −5 on a Vitrobot Mark IV (FEI) at room temperature and 100% humidity, then plunged into liquid ethane. For the second dataset, the final Pol I concentration in the stalled EC sample was 0.24 mg/ml and no further chemicals were added. 4 µl of sample were applied to freshly glow-discharged Quantifoil R2/2, 300 mesh grids. After 30 s incubation, grids were blotted for 3 s and offset –3 mm on a Vitrobot Mark II (FEI) at room temperature and 100% humidity, then plunged into liquid ethane. In both cases, vitrified grids were stored in liquid nitrogen. Both datasets were collected on the Titan Krios I electron microscope (FEI) at the Diamond Light Source operated at 300 kV, using a K2 summit direct electron detector (Gatan) and the EPU automated single-particle acquisition software (FEI). In both cases, data were acquired with a final pixel size of 1.047 Å at the specimen level, using defocus values between –1 and –3.5 µm in 0.3 µm intervals, and the camera was operated in dose-fractionation counting mode. For grids coated with thin continuous carbon, 4425 movies were collected at tilt angle 0°, with a total dose of 42.5 e^-^/Å^2^ accumulated over 8 s and fractionated across 40 frames. For holey grids lacking continuous carbon, 6424 movies were collected at tilt angles 0°, 20° and 30° with a total dose of 40.1 e^-^/Å^2^ accumulated over 8 s and fractionated across 40 frames.

For the sample containing RNA-2, the final Pol I concentration was 0.4 mg/ml in the presence of 0.5 mM EDTA. 4 µl of sample were applied to freshly glow-discharged UltrAufoil R1.2/1.3, 300 mesh grids. After 30 s incubation, grids were blotted for 2 s and blot force of −5 on a Vitrobot Mark IV (FEI) at 10 °C and 100% humidity, then plunged into liquid ethane and stored in liquid nitrogen. Data were collected on the Titan Krios (FEI) at the BREM Facility operated at 300 kV, using a K3 direct electron detector (Gatan) and the EPU automated single-particle acquisition software (FEI). Data were acquired with a final pixel size of 0.646 Å at the specimen level, using defocus values between –1 and –2.5 µm in 0.3 µm intervals, and the camera was operated in dose-fractionation counting mode. A total of 23488 movies were collected at tilt angle 30° with a total dose of 47.2 e^-^/Å^2^ accumulated over 1.7 s and fractionated across 40 frames.

### Cryo-EM data processing

For the sample containing RNA-1, a total of 10849 movies from both datasets were aligned with MotionCorr2^59^ and global CTF parameters were estimated using CtfFind^60^. 2137 of those were discarded after manual inspection of average micrographs and their corresponding power spectra. For the first dataset, around 1000 particles were manually-picked, reference-free 2D classified and the resulting averages used as templates for autopicking in Relion 3.0^61^. For the second dataset, particle locations were obtained with crYOLO^62^, then used to estimate per-particle CTF defocus values with Gctf^63^. Relion 3.0 was used for further processing. Particles from both datasets were separately extracted using a box of 288 pixels and subjected to several rounds of 2D classification. 199450 and 635120 particles from the first and second datasets, respectively, belonging to classes that produce averages showing secondary structure features were joined and used to generate an initial model. All particles were jointly refined using the initial model as a reference, then polished and their CTF parameters refined. A global 3D classification into 6 classes was performed. 2 classes lacking two or three lobe-associated subunits, plus 2 classes with different cleft opening were refined. A fifth class showed density in the funnel and was used for focused 3D classification using a mask around the funnel. The resulting class showing strong density in the funnel was subsequently refined. To identify AMPCPP binding sites, we performed focused 3D classification with a mask around the active center using CryoSPARC^64^. Two classes showing density accounting for AMPCPP at distinct locations in the vicinity of the active center were refined. In all cases, final post-processing was performed using automatic masking and B-factor sharpening.

For the sample containing RNA-2, a total of 23488 movies were imported into CryoSPARC, where subsequent image analysis was performed. After movie alignment and CTF estimation, 4234 movies were discarded. Particles were selected using the blob picking utility, then extracted using a box of 500 pixels and subjected to several rounds of 2D classification. 1282765 particles belonging to classes with averages showing secondary structure features were used to generate an ab initio model. These particles were subjected to focused 3D classification with a mask around the DNA-binding cleft. One of the resulting classes showing density for the RNA 3’-end was refined using non-uniform refinement.

### Model building and refinement

The available structure of undamaged Pol I EC containing GMPCPP in the A-site (PDB code: 6HKO) was fitted into the 3D maps using UCSF Chimera^65^ and used as starting point for model building with Coot^66^ and subsequent real-space refinement as implemented in Phenix^67^. Refinement statistics are summarized in Table S1. Figures were prepared using PyMOL (Schrödinger Inc.) and ChimeraX^68^. Sequence alignments are performed with Clustalw^69^ and represented with ESPript3^70^.

### Data availability

The structures described here and their associated data have been deposited in the Protein Data Bank under accession codes 9G1V, 9G26, 9G2C, 9G29, 9G2B, 9G1X, 9G23, 9G24, 9G27, and the Electron Microscopy Data Bank under accession codes 50955, 50965, 50972, 50970, 50971, 50956, 50962, 50963, 50966, as described in Table S1.

## Acknowledgements

We thank J. Varela and C. Quevedo at the Protein Chemistry Facility (CIB-CSIC) and R. Núñez at the Electron Microscopy Facility (CIB-CSIC) for technical assistance. We acknowledge the Diamond Light Source for access and support of the cryo-EM facilities at the UK national electron Bio-Imaging Centre (eBIC). We also thank the Basque Resource for Electron Microscopy (BREM) located at Instituto Biofisika (UPV/EHU, CSIC), supported by the Department of Education and the Innovation Fund of the Basque Government, Fundación Biofísica Bizkaia and MCIN with funding from European Union NextGenerationEU (PRTR-C17.I1). This work was supported by the Spanish Ministry of Science / Agencia Estatal de Investigación (PID2020-116722GB-I00 to C.F.-T.; PRE2021-098362 to A.S.-A.; PRE2018-087012 to A.P.-P.) and intramural funding from CSIC (2020AEP152 and PIE-202120E047-Conexiones-Life to C.F.-T.). A.S.-A. acknowledges support from the Madrid City Council for a living stipend at Residencia de Estudiantes.

## References

1. Lindahl, T. (1993). Instability and decay of the primary structure of DNA. Nature 362, 709–715. 10.1038/362709a0.

2. Loeb, L.A., and Preston, B.D. (1986). Mutagenesis by apurinic/apyrimidinic sites. Annu. Rev. Genet. 20, 201–230. 10.1146/annurev.ge.20.120186.001221.

3. Strauss, B.S. (1991). The ‘A rule’ of mutagen specificity: A consequence of DNA polymerase bypass of non-instructional lesions? BioEssays 13, 79–84. 10.1002/bies.950130206.

4. Zahn, K.E., Belrhali, H., Wallace, S.S., and Doublié, S. (2007). Caught Bending the A-Rule: Crystal Structures of Translesion DNA Synthesis with a Non-Natural Nucleotide. Biochemistry 46, 10551–10561. 10.1021/bi7008807.

5. Obeid, S., Blatter, N., Kranaster, R., Schnur, A., Diederichs, K., Welte, W., and Marx, A. (2010). Replication through an abasic DNA lesion: structural basis for adenine selectivity. EMBO J. 29, 1738–1747. 10.1038/emboj.2010.64.

6. Obeid, S., Welte, W., Diederichs, K., and Marx, A. (2012). Amino Acid Templating Mechanisms in Selection of Nucleotides Opposite Abasic Sites by a Family A DNA Polymerase*. J. Biol. Chem. 287, 14099–14108. 10.1074/jbc.M111.334904.

7. Beard, W.A., Shock, D.D., Batra, V.K., Pedersen, L.C., and Wilson, S.H. (2009). DNA Polymerase β Substrate Specificity. J. Biol. Chem. 284, 31680–31689. 10.1074/jbc.M109.029843.

8. Nair, D.T., Johnson, R.E., Prakash, L., Prakash, S., and Aggarwal, A.K. (2009). DNA Synthesis across an Abasic Lesion by Human DNA Polymerase ι. Structure 17, 530–537. 10.1016/j.str.2009.02.015.

9. Wang, W., Walmacq, C., Chong, J., Kashlev, M., and Wang, D. (2018). Structural basis of transcriptional stalling and bypass of abasic DNA lesion by RNA polymerase II. Proc. Natl. Acad. Sci. 115, E2538–E2545. 10.1073/pnas.1722050115.

10. Gnatt, A.L., Cramer, P., Fu, J., Bushnell, D.A., and Kornberg, R.D. (2001). Structural basis of transcription: an RNA polymerase II elongation complex at 3.3 A resolution. Science 292, 1876–1882. 10.1126/science.1059495.

11. Kettenberger, H., Armache, K.-J., and Cramer, P. (2004). Complete RNA polymerase II elongation complex structure and its interactions with NTP and TFIIS. Mol. Cell 16, 955–965. 10.1016/j.molcel.2004.11.040.

12. Wang, D., Bushnell, D.A., Westover, K.D., Kaplan, C.D., and Kornberg, R.D. (2006). Structural basis of transcription: role of the trigger loop in substrate specificity and catalysis. Cell 127, 941–954. 10.1016/j.cell.2006.11.023.

13. Cheung, A.C.M., and Cramer, P. (2011). Structural basis of RNA polymerase II backtracking, arrest and reactivation. Nature 471, 249–253. 10.1038/nature09785.

14. Brueckner, F., and Cramer, P. (2008). Structural basis of transcription inhibition by α-amanitin and implications for RNA polymerase II translocation. Nat. Struct. Mol. Biol. 15, 811–818. 10.1038/nsmb.1458.

15. Westover, K.D., Bushnell, D.A., and Kornberg, R.D. (2004). Structural Basis of Transcription: Nucleotide Selection by Rotation in the RNA Polymerase II Active Center. Cell 119, 481–489. 10.1016/j.cell.2004.10.016.

16. Landick, R. (2006). The regulatory roles and mechanism of transcriptional pausing. Biochem. Soc. Trans. 34, 1062– 1066. 10.1042/BST0341062.

17. Wang, D., Bushnell, D.A., Huang, X., Westover, K.D., Levitt, M., and Kornberg, R.D. (2009). Structural Basis of Transcription: Backtracked RNA Polymerase II at 3.4 Angstrom Resolution. Science 324, 1203–1206. 10.1126/science.1168729.

18. Warner, J.R. (1999). The economics of ribosome biosynthesis in yeast. Trends Biochem. Sci. 24, 437–440. 10.1016/S0968-0004(99)01460-7.

19. Engel, C., Sainsbury, S., Cheung, A.C., Kostrewa, D., and Cramer, P. (2013). RNA polymerase I structure and transcription regulation. Nature 502, 650–655. 10.1038/nature12712.

20. Fernández-Tornero, C., Moreno-Morcillo, M., Rashid, U.J., Taylor, N.M.I., Ruiz, F.M., Gruene, T., Legrand, P., Steuerwald, U., and Müller, C.W. (2013). Crystal structure of the 14-subunit RNA polymerase I. Nature 502, 644– 649. 10.1038/nature12636.

21. Engel, C., Plitzko, J., and Cramer, P. (2016). RNA polymerase I–Rrn3 complex at 4.8 Å resolution. Nat. Commun. 7, 12129. 10.1038/ncomms12129.

22. Pilsl, M., Crucifix, C., Papai, G., Krupp, F., Steinbauer, R., Griesenbeck, J., Milkereit, P., Tschochner, H., and Schultz, P. (2016). Structure of the initiation-competent RNA polymerase I and its implication for transcription. Nat. Commun. 7, 12126. 10.1038/ncomms12126.

23. Torreira, E., Louro, J.A., Pazos, I., González-Polo, N., Gil-Carton, D., Duran, A.G., Tosi, S., Gallego, O., Calvo, O., and Fernández-Tornero, C. (2017). The dynamic assembly of distinct RNA polymerase I complexes modulates rDNA transcription. eLife 6, e20832. 10.7554/eLife.20832.

24. Yamamoto, R.T., Nogi, Y., Dodd, J.A., and Nomura, M. (1996). RRN3 gene of Saccharomyces cerevisiae encodes an essential RNA polymerase I transcription factor which interacts with the polymerase independently of DNA template. EMBO J. 15, 3964–3973.

25. Keener, J., Josaitis, C.A., Dodd, J.A., and Nomura, M. (1998). Reconstitution of Yeast RNA Polymerase I Transcription *in Vitro* from Purified Components. J. Biol. Chem. 273, 33795–33802. 10.1074/jbc.273.50.33795.

26. 26. Van Mullem, V., Landrieux, E., Vandenhaute, J., and Thuriaux, P. (2002). Rpa12p, a conserved RNA polymerase I subunit with two functional domains. Mol. Microbiol. 43, 1105–1113. 10.1046/j.1365-2958.2002.02824.x.

27. Appling, F.D., Schneider, D.A., and Lucius, A.L. (2017). Multisubunit RNA Polymerase Cleavage Factors Modulate the Kinetics and Energetics of Nucleotide Incorporation: An RNA Polymerase I Case Study. Biochemistry 56, 5654–5662. 10.1021/acs.biochem.7b00370.

28. Scull, C.E., Lucius, A.L., and Schneider, D.A. (2021). The N-terminal domain of the A12.2 subunit stimulates RNA polymerase I transcription elongation. Biophys. J. 120, 1883–1893. 10.1016/j.bpj.2021.03.007.

29. Kuhn, C.-D., Geiger, S.R., Baumli, S., Gartmann, M., Gerber, J., Jennebach, S., Mielke, T., Tschochner, H., Beckmann, R., and Cramer, P. (2007). Functional Architecture of RNA Polymerase I. Cell 131, 1260–1272. 10.1016/j.cell.2007.10.051.

30. Schwank, K., Schmid, C., Fremter, T., Milkereit, P., Griesenbeck, J., and Tschochner, H. (2022). RNA polymerase I (Pol I) lobe-binding subunit Rpa12.2 promotes RNA cleavage and proofreading. J. Biol. Chem. 298, 101862. 10.1016/j.jbc.2022.101862.

31. Geiger, S.R., Lorenzen, K., Schreieck, A., Hanecker, P., Kostrewa, D., Heck, A.J.R., and Cramer, P. (2010). RNA polymerase I contains a TFIIF-related DNA-binding subcomplex. Mol. Cell 39, 583–594. 10.1016/j.molcel.2010.07.028.

32. Huet, J., Buhler, J.M., Sentenac, A., and Fromageot, P. (1975). Dissociation of two polypeptide chains from yeast RNA polymerase A. Proc. Natl. Acad. Sci. 72, 3034–3038. 10.1073/pnas.72.8.3034.

33. Tafur, L., Sadian, Y., Hanske, J., Wetzel, R., Weis, F., and Müller, C.W. (2019). The cryo-EM structure of a 12-subunit variant of RNA polymerase I reveals dissociation of the A49-A34.5 heterodimer and rearrangement of subunit A12.2. eLife 8, e43204. 10.7554/eLife.43204.

34. Sanz-Murillo, M., Xu, J., Belogurov, G.A., Calvo, O., Gil-Carton, D., Moreno-Morcillo, M., Wang, D., and Fernández-Tornero, C. (2018). Structural basis of RNA polymerase I stalling at UV light-induced DNA damage. Proc. Natl. Acad. Sci. U. S. A. 115, 8972–8977. 10.1073/pnas.1802626115.

35. Tornaletti, S., Maeda, L.S., and Hanawalt, P.C. (2006). Transcription Arrest at an Abasic Site in the Transcribed Strand of Template DNA. Chem. Res. Toxicol. 19, 1215–1220. 10.1021/tx060103g.

36. Neyer, S., Kunz, M., Geiss, C., Hantsche, M., Hodirnau, V.-V., Seybert, A., Engel, C., Scheffer, M.P., Cramer, P., and Frangakis, A.S. (2016). Structure of RNA polymerase I transcribing ribosomal DNA genes. Nature 540, 607– 610. 10.1038/nature20561.

37. Sadian, Y., Baudin, F., Tafur, L., Murciano, B., Wetzel, R., Weis, F., and Müller, C.W. (2019). Molecular insight into RNA polymerase I promoter recognition and promoter melting. Nat. Commun. 10, 5543. 10.1038/s41467-019-13510-w.

38. Nguyen, P.Q., Huecas, S., Asif-Laidin, A., Plaza-Pegueroles, A., Capuzzi, B., Palmic, N., Conesa, C., Acker, J., Reguera, J., Lesage, P., et al. (2023). Structural basis of Ty1 integrase tethering to RNA polymerase III for targeted retrotransposon integration. Nat. Commun. 14, 1729. 10.1038/s41467-023-37109-4.

39. Fernández-Tornero, C. (2018). RNA polymerase I activation and hibernation: unique mechanisms for unique genes. Transcription 9, 248–254. 10.1080/21541264.2017.1416267.

40. Mccusker, J.H., Yamagish, M., Kolb, J.M., and Nomura, M. (1991). Suppressor Analysis of Temperature-Sensitive RNA Polymerase I Mutations in Saccharomyces cerevisiae: Suppression of Mutations in a Zinc-Binding Motif by Transposed Mutant Genes. Mol. Cell. Biol. 11, 746–753. 10.1128/mcb.11.2.746-753.1991.

41. Gadal, O., Mariotte-Labarre, S., Chedin, S., Quemeneur, E., Carles, C., Sentenac, A., and Thuriaux, P. (1997). A34.5, a Nonessential Component of Yeast RNA Polymerase I, Cooperates with Subunit A14 and DNA Topoisomerase I To Produce a Functional rRNA Synthesis Machine†. Mol. Cell. Biol. 17, 1787–1795. 10.1128/MCB.17.4.1787.

42. Kang, J.Y., Mishanina, T.V., Bellecourt, M.J., Mooney, R.A., Darst, S.A., and Landick, R. (2018). RNA Polymerase Accommodates a Pause RNA Hairpin by Global Conformational Rearrangements that Prolong Pausing. Mol. Cell 69, 802–815.e5. 10.1016/j.molcel.2018.01.018.

43. Su, B.G., and Vos, S.M. (2024). Distinct negative elongation factor conformations regulate RNA polymerase II promoter-proximal pausing. Mol. Cell 84, 1243–1256.e5. 10.1016/j.molcel.2024.01.023.

44. Jacobs, R.Q., Carter, Z.I., Lucius, A.L., and Schneider, D.A. (2022). Uncovering the mechanisms of transcription elongation by eukaryotic RNA polymerases I, II, and III. iScience 25, 105306. 10.1016/j.isci.2022.105306.

45. Batada, N.N., Westover, K.D., Bushnell, D.A., Levitt, M., and Kornberg, R.D. (2004). Diffusion of nucleoside triphosphates and role of the entry site to the RNA polymerase II active center. Proc. Natl. Acad. Sci. 101, 17361– 17364. 10.1073/pnas.0408168101.

46. Lisica, A., Engel, C., Jahnel, M., Roldán, É., Galburt, E.A., Cramer, P., and Grill, S.W. (2016). Mechanisms of backtrack recovery by RNA polymerases I and II. Proc. Natl. Acad. Sci. 113, 2946–2951. 10.1073/pnas.1517011113.

47. Schwank, K., Schmid, C., Fremter, T., Engel, C., Milkereit, P., Griesenbeck, J., and Tschochner, H. (2023). Features of yeast RNA polymerase I with special consideration of the lobe binding subunits. Biol. Chem. 404, 979–1002. 10.1515/hsz-2023-0184.

48. Liljelund, P., Mariotte, S., Buhler, J.M., and Sentenac, A. (1992). Characterization and mutagenesis of the gene encoding the A49 subunit of RNA polymerase A in Saccharomyces cerevisiae. Proc. Natl. Acad. Sci. 89, 9302– 9305. 10.1073/pnas.89.19.9302.

49. Beckouet, F., Labarre-Mariotte, S., Albert, B., Imazawa, Y., Werner, M., Gadal, O., Nogi, Y., and Thuriaux, P. (2008). Two RNA Polymerase I Subunits Control the Binding and Release of Rrn3 during Transcription. Mol. Cell. Biol. 28, 1596–1605. 10.1128/MCB.01464-07.

50. Rossi, M.J., Kuntala, P.K., Lai, W.K.M., Yamada, N., Badjatia, N., Mittal, C., Kuzu, G., Bocklund, K., Farrell, N.P., Blanda, T.R., et al. (2021). A high-resolution protein architecture of the budding yeast genome. Nature 592, 309–314. 10.1038/s41586-021-03314-8.

51. Farnung, L., Ochmann, M., Engeholm, M., and Cramer, P. (2021). Structural basis of nucleosome transcription mediated by Chd1 and FACT. Nat. Struct. Mol. Biol. 28, 382–387. 10.1038/s41594-021-00578-6.

52. Viktorovskaya, O.V., Appling, F.D., and Schneider, D.A. (2011). Yeast Transcription Elongation Factor Spt5 Associates with RNA Polymerase I and RNA Polymerase II Directly*. J. Biol. Chem. 286, 18825–18833. 10.1074/jbc.M110.202119.

53. Appling, F.D., Scull, C.E., Lucius, A.L., and Schneider, D.A. (2018). The A12.2 Subunit Is an Intrinsic Destabilizer of the RNA Polymerase I Elongation Complex. Biophys. J. 114, 2507–2515. 10.1016/j.bpj.2018.04.015.

54. Kim, N., and Jinks-Robertson, S. (2010). Abasic Sites in the Transcribed Strand of Yeast DNA Are Removed by Transcription-Coupled Nucleotide Excision Repair. Mol. Cell. Biol. 30, 3206–3215. 10.1128/MCB.00308-10.

55. Swanson, R.L., Morey, N.J., Doetsch, P.W., and Jinks-Robertson, S. (1999). Overlapping Specificities of Base Excision Repair, Nucleotide Excision Repair, Recombination, and Translesion Synthesis Pathways for DNA Base Damage in Saccharomyces cerevisiae. Mol. Cell. Biol. 19, 2929–2935. 10.1128/MCB.19.4.2929.

56. Torres-Ramos, C.A., Johnson, R.E., Prakash, L., and Prakash, S. (2000). Evidence for the Involvement of Nucleotide Excision Repair in the Removal of Abasic Sites in Yeast. Mol. Cell. Biol. 20, 3522–3528. 10.1128/MCB.20.10.3522-3528.2000.

57. Hanawalt, P.C., and Spivak, G. (2008). Transcription-coupled DNA repair: two decades of progress and surprises. Nat. Rev. Mol. Cell Biol. 9, 958–970. 10.1038/nrm2549.

58. Moreno-Morcillo, M., Taylor, N.M.I., Gruene, T., Legrand, P., Rashid, U.J., Ruiz, F.M., Steuerwald, U., Müller, C.W., and Fernández-Tornero, C. (2014). Solving the RNA polymerase I structural puzzle. Acta Crystallogr. D Biol. Crystallogr. 70, 2570–2582. 10.1107/S1399004714015788.

59. Zheng, S.Q., Palovcak, E., Armache, J.-P., Verba, K.A., Cheng, Y., and Agard, D.A. (2017). MotionCor2: anisotropic correction of beam-induced motion for improved cryo-electron microscopy. Nat. Methods 14, 331–332. 10.1038/nmeth.4193.

60. Rohou, A., and Grigorieff, N. (2015). CTFFIND4: Fast and accurate defocus estimation from electron micrographs. J. Struct. Biol. 192, 216–221. 10.1016/j.jsb.2015.08.008.

61. Zivanov, J., Nakane, T., Forsberg, B.O., Kimanius, D., Hagen, W.J., Lindahl, E., and Scheres, S.H. (2018). New tools for automated high-resolution cryo-EM structure determination in RELION-3. eLife 7, e42166. 10.7554/eLife.42166.

62. Wagner, T., Merino, F., Stabrin, M., Moriya, T., Antoni, C., Apelbaum, A., Hagel, P., Sitsel, O., Raisch, T., Prumbaum, D., et al. (2019). SPHIRE-crYOLO is a fast and accurate fully automated particle picker for cryo-EM. Commun. Biol. 2, 1–13. 10.1038/s42003-019-0437-z.

63. Zhang, K. (2016). Gctf: Real-time CTF determination and correction. J. Struct. Biol. 193, 1–12. 10.1016/j.jsb.2015.11.003.

64. Punjani, A., Rubinstein, J.L., Fleet, D.J., and Brubaker, M.A. (2017). cryoSPARC: algorithms for rapid unsupervised cryo-EM structure determination. Nat. Methods 14, 290–296. 10.1038/nmeth.4169.

65. Pettersen, E.F., Goddard, T.D., Huang, C.C., Couch, G.S., Greenblatt, D.M., Meng, E.C., and Ferrin, T.E. (2004). UCSF Chimera—A visualization system for exploratory research and analysis. J. Comput. Chem. 25, 1605–1612. 10.1002/jcc.20084.

66. Emsley, P., Lohkamp, B., Scott, W.G., and Cowtan, K. (2010). Features and development of Coot. Acta Crystallogr. D Biol. Crystallogr. 66, 486–501. 10.1107/S0907444910007493.

67. Liebschner, D., Afonine, P.V., Baker, M.L., Bunkóczi, G., Chen, V.B., Croll, T.I., Hintze, B., Hung, L.-W., Jain, S., McCoy, A.J., et al. (2019). Macromolecular structure determination using X-rays, neutrons and electrons: recent developments in Phenix. Acta Crystallogr. Sect. Struct. Biol. 75, 861–877. 10.1107/S2059798319011471.

68. Meng, E.C., Goddard, T.D., Pettersen, E.F., Couch, G.S., Pearson, Z.J., Morris, J.H., and Ferrin, T.E. (2023). UCSF ChimeraX: Tools for structure building and analysis. Protein Sci. 32, e4792. 10.1002/pro.4792.

69. Sievers, F., and Higgins, D.G. (2018). Clustal Omega for making accurate alignments of many protein sequences. Protein Sci. Publ. Protein Soc. 27, 135–145. 10.1002/pro.3290.

70. Robert, X., and Gouet, P. (2014). Deciphering key features in protein structures with the new ENDscript server. Nucleic Acids Res. 42, W320–324. 10.1093/nar/gku316.

